# Predicting the effects of drug combinations using probabilistic matrix factorization

**DOI:** 10.1101/2021.05.03.442470

**Authors:** Ron Nafshi, Timothy R. Lezon

## Abstract

Drug development is costly and time-consuming, and developing novel practical strategies for creating more effective treatments is imperative. One possible solution is to prescribe drugs in combination. Synergistic drug combinations could allow lower doses of each constituent drug, reducing adverse reactions and drug resistance. However, it is not feasible to sufficiently test every combination of drugs for a given illness to determine promising synergistic combinations. Since there is a finite amount of time and resources available for finding synergistic combinations, a model that can identify synergistic combinations from a limited subset of all available combinations could accelerate development of therapeutics. By applying recommender algorithms, such as the low-rank matrix completion algorithm Probabilistic Matrix Factorization (PMF), it may be possible to identify synergistic combinations from partial information of the drug interactions. Here, we use PMF to predict the efficacy of two-drug combinations using the NCI ALMANAC, a robust collection of pairwise drug combinations of 104 FDA-approved anticancer drugs against 60 common cancer cell lines. We find that PMF is able predict drug combination efficacy with high accuracy from a limited set of combinations and is robust to changes in the individual training data. Moreover, we propose a new PMF-guided experimental design to detect all synergistic combinations without testing every combination.

## Introduction

Despite recent advances in drug development and disease biology, the cost to develop a new effective drug remains prohibitively high; as of 2019, the estimated cost to develop new prescription medicine that gains marketing approval is estimated to be $2.6 billion. Furthermore, the approval rate of drugs in clinical testing is an abysmal 12 percent [2]. Thus, developing practical strategies for creating more effective treatments is imperative. One possible solution is to prescribe drugs in combination. The use of synergistic drug combinations has several major benefits: it can reduce development of drug resistance, and it can allow for lower dosages of each constituent drug, lessening the adverse effects of each [3]. However, it is simply not feasible to sufficiently test every combination of drugs for a given illness to determine promising synergistic combinations. Since there is a limited amount of time and resources available for testing of synergistic combinations, a model that can identify synergistic combinations from a limited subset of all available combinations could accelerate development of therapies. By using collaborative filtering algorithms, such as Probabilistic Matrix Factorization (PMF) [4], it may be possible to identify synergistic combinations from partial information of drug-drug interactions.

Imputing missing data values is a longstanding problem that has been addressed in a variety of ways, with algorithms such as pairwise deletion, mean substitution, and k-nearest neighbor. Pairwise deletion and mean substitution work efficiently in small datasets and are comparatively fast, but have several drawbacks, such as losing data, biasing the sample statistics, and not accounting for the correlation between features [5]. K-Nearest Neighbor, while simple, has several limitations, such as not handling sparsity well and being computationally expensive as the input size grows. PMF has the advantage that its computation time scales linearly and it can make accurate predictions for sparse and imbalanced data sets [4].

The PMF algorithm was developed to recommend movies to Netflix users based on the movies viewed by other users. Given that no Netflix user rated every movie, the core assumption of PMF is that attitudes or preferences that lead to each user’s score for a movie are based on a small number of unobserved factors. Thus, PMF models each user’s recommendations as a linear combination of item factor vectors using user-specific coefficients [4]. The method has since been applied to predict values from other large, sparse and imbalanced data sets. Biomedical applications of PMF include predicting diseases associated with transcription patterns [6, 7], recommending novel indications for drug repurposing [8, 9], and predicting novel targets from drugs [10–12]. Here, we use PMF to predict the effects of novel two-drug combinations based on information from other two-drug combinations. We train our model on phenotypic screening data from the NCI ALMANAC [1], a robust collection of pairwise drug combinations of 104 FDA approved anticancer drugs against 60 common cancer cell lines. We find that knowing the effects of only 70% of drug combinations allows us to classify the effects of the missing combinations as efficacious with 95% accuracy, and we demonstrate how our method can be incorporated into optimal experimental design.

## Methods

### NCI ALMANAC

The NCI ALMANAC is a novel, easy-to-use resource created to help researchers identify new combination therapies. The NCI ALMANAC database [1] is a collection of pairwise combinations of 104 FDA approved anticancer drugs against the NCI-60, a set of 60 common human tumor cancer cell lines collected by the National Cancer Institute. A total of 5,232 drug-drug pairs were evaluated in each of the cell lines; 304,549 experiments were performed to test each drug at either 9 or 15 combination dose points, for a total of 2,809,671 dose combinations. At each dose combination, the percent growth after two days was measured and recorded, and combination efficacy derived. The synergy of each combination is reported by a “ComboScore” that measures the difference between the recorded growth rate after testing and the growth rate expected by Bliss Independence[13]. A positive ComboScore indicates a synergistic combination, whereas a negative ComboScore indicates an antagonistic combination. For each cell line, the ComboScores and combination efficacies are arranged into a symmetric matrix, **M**_104*x*104_, where each element represents the ComboScore or efficacy of a unique drug-drug combination on that cell line. For purposes of PMF, diagonal elements are ignored. The data is then mean-zero standardized for input into the PMF algorithm.

### PMF

Probabilistic Matrix Factorization (PMF) is a collaborative filtering algorithm that factors the low-rank input matrix **M**_*n×m*_ into the product of two low-rank matrices, **A**_*n×d*_ and **B**_*m×d*_ such that 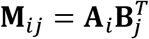. Thus, PMF reduces to estimating the two matrices **A** and **B**. The core assumptions of this are that the values of **M** are independent, normally distributed and share a common variance *σ*^2^. Thus the conditional probability of entries of **M** can be expressed as: 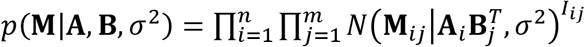, where *I_ij_* is the indicator function equal to 1 if **M**_*ij*_ is known and 0 otherwise [4].

To solve for the matrices **A** and **B**, we use gradient descent with gradient acceleration. Stochastic gradient descent methods are a critical component of machine learning, and methods incorporating momentum and acceleration play an important role when used in conjunction with stochastic gradients [14]. Momentum methods help accelerate stochastic gradient descent in the relevant direction and dampen oscillations as a minimum is approached by incorporating the momentum constant *γ*. The update step with respect to the parameters *θ* can be expressed as *ν_t_* = *γν*_*t*−1_ + *η*∇_*θ*_*J*/(*θ*), *θ* = *θ* – *ν_t_*. However, simple momentum methods can be insufficient for complex surfaces. The Nesterov Accelerated Gradient (NAG) [14] improves on this method by “looking ahead” to where the parameters will be to calculate the gradient and is formalized as followed: *ν_t_* = *γν*_*t*−1_ + *η*∇_*θ*_J(*θ* – *γν*_*t*−1_), *θ* = *θ* – *ν_t_*. Rather than computing the gradient at parameters *θ*, NAG looks ahead at a rough approximation of where the parameters will be, computing the gradient at *θ* – *γν*_*t*−1_. This anticipatory update greatly increases optimization and performance of PMF as it approaches a minimum.

## Results and Discussion

### PMF accurately recovers drug synergies from partial data

We first investigated the ability of PMF to recover hidden elements in the drug combination efficacy matrix. For each cell line, we randomly hid a fraction of the combination efficacy matrix and used PMF to predict the hidden values. To guarantee a solution, we included only cases where all drugs were present in a single connected component; that is, where a path could be made from any drug to any other drug using common combination partners. PMF recovered training data to arbitrary precision (**Fig. 1a**) and recovered test data well, provided a sufficiently large training set (i.e., small fraction of data hidden). Using empirically determined hyperparameters, we found that knowing only 30-50% of the drug-drug interactions was sufficient to recover the remaining values in the matrix to within 10% (**Fig. 1b,c**).

**Figure 1.**
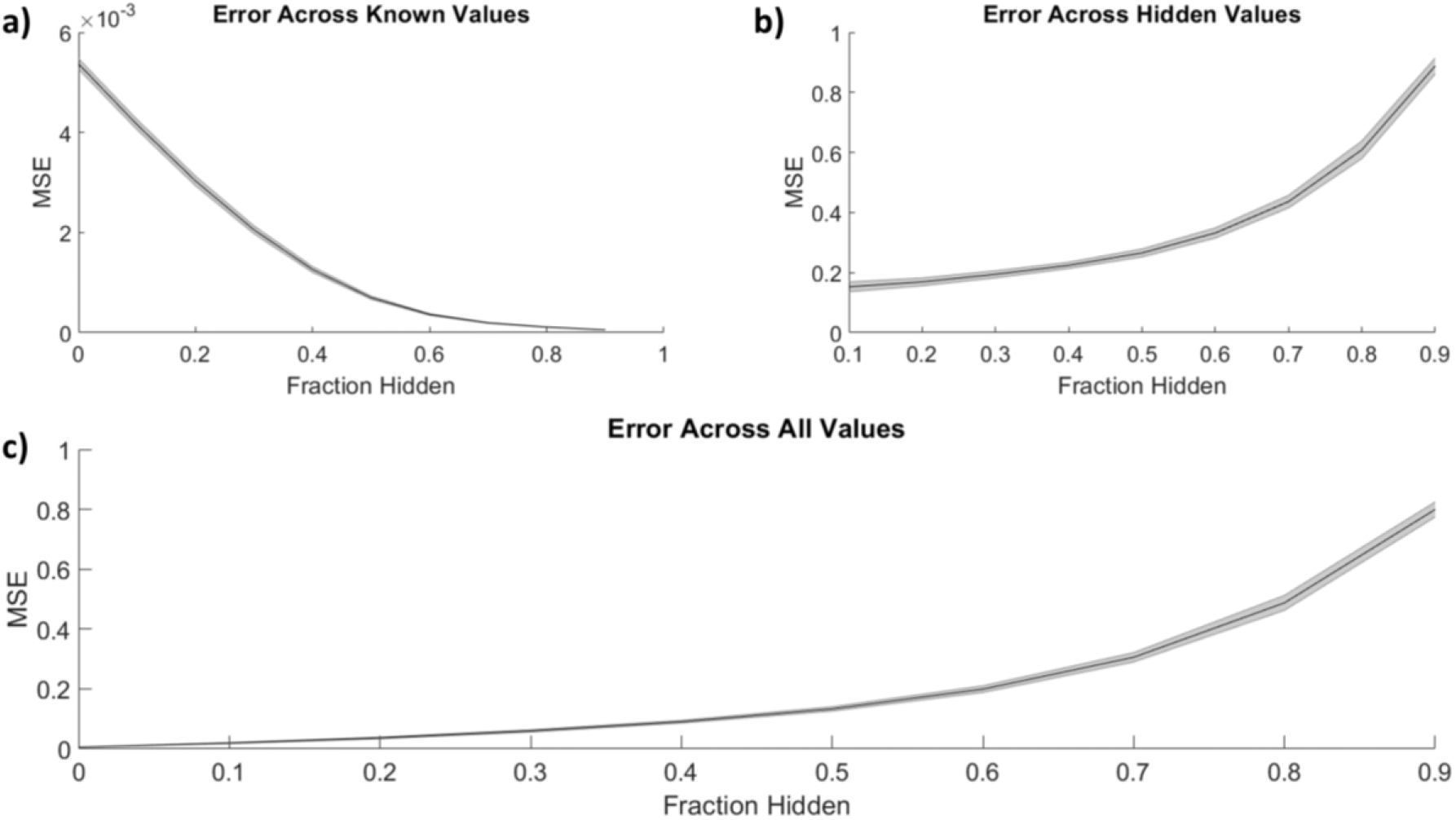
PMF recovers the values of hidden elements of the drug efficacy matrix from only a fraction of interactions. The mean-squared error of PMF in recovering values of a) known, b) hidden, and c) all elements is plotted against the fraction of hidden data. In all panels, the shaded area represents the standard deviation of the mean-squared error over 25 trials across all cell lines.

When selecting combinations with efficacies above a given threshold, PMF performance did not vary strongly with the threshold value (**Fig. 2**); that is, the method can predict whether a combination has an effect over 0.9 nearly as well as it can predict whether a combination has an effect over 0.2.

**Figure 2.**
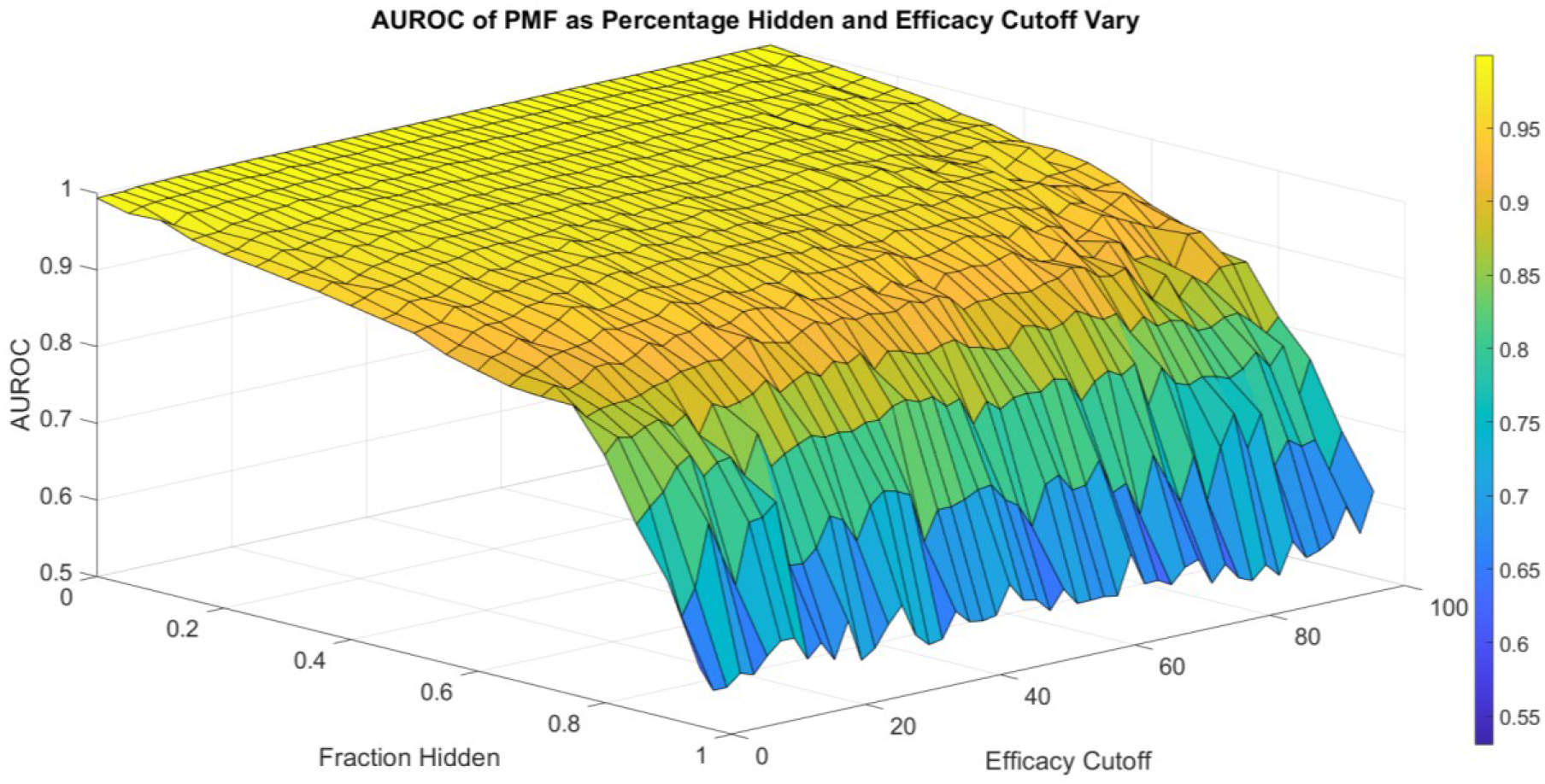
The area under the ROC curve (AUROC) of PMF is shown as the fraction hidden and efficacy cutoff vary on the 786-0 cell line, which is representative of all cell lines. The efficacy cutoff describes the efficacy at which a drug-drug combination is considered active, with combination efficacy defined as 100 minus the percent growth as described in the standard NCI-60 testing protocol [1]. As the fraction hidden decreases, the performance of the model remains high until it drops sharply at 70% hidden and performs with similar accuracy regardless of the efficacy cutoff, decaying to random guesses when the full matrix is hidden. The smooth surface indicates PMF reproduces all elements with equal accuracy and is not heavily affected by outliers.

### PMF performance is largely independent of individual drug efficacies

Assuming compounds act independently (i.e., Bliss independence), the most efficacious compound combinations will be combinations of the independently most efficacious compounds. Reasoning that efficacious drugs are more likely to influence pathologically relevant mechanisms, we next investigated whether PMF performed better when trained on combinations involving highly efficacious drugs. For each cell line, we rank-ordered the compounds by efficacy and compared the accuracy of PMF using only the top 52 individually efficacious drugs and PMF using only the bottom 52 individually efficacious drugs.

PMF was slightly more accurate at predicting smaller subsets of the drug-efficacy matrix using only highly efficacious drugs rather than weakly efficacious combinations (**Suppl. Fig. 1**). However, on aggregate the differences were small, and PMF performance was largely independent of the individual efficacies of the starting set outside of this edge case. More generally, we found that the most efficacious compounds neither led to the most efficacious combinations, nor were they the best at predicting the values of missing efficacies (**Suppl. Fig. 1**). In fact, individual drug identities did not greatly affect the accuracy of the prediction. We generated an occupancy matrix by randomly selecting 10% of the elements in the combination efficacy matrix. We then randomly shuffled the identities of the drugs while keeping the occupancy matrix static. Repeating this 1000 times for 1000 different occupancy matrices, we found PMF predicted the missing values of each matrix with a mean squared error of 0.938+/0.0145, and thus performed equally well regardless of the individual drug identities for a given occupancy matrix.

### Graph topology’s influence on PMF performance

The combination efficacy matrix describes an undirected graph in which the *N* drugs are nodes and the edges represent two-drug combinations. The challenge of PMF is to reconstruct the fully connected graph from a seed network. By using different algorithms for selecting drug combinations for the training set, we investigated how seed network topology influences prediction accuracy. The method described above, where seed drug combinations are selected randomly and independently, results in an Erdős-Réyni graph [15] that has a Poisson degree distribution [16].

An extension of the Erdős-Réyni graph is the Watts-Strogatz model [17]. This method is motivated by the observation that real networks often have the Small-World Property [18] and high clustering, and it generates random distributions following these ideas. The Watts-Strogatz graph is generated by attaching each node to its nearest *k* neighbors, resulting in a regular lattice structure. Each edge is then randomly reassigned with probability β. When β is 0, no changes are accepted, and the method preserves the original lattice. As β increases, more links will be randomly assigned, and as β approaches 1, all links will be reassigned, resulting in a completely random Erdős-Réyni network. Intermediate values of β result in small-world networks of low diameter [16].

When training data was arranged in a Watts-Strogatz model topology, the performance of PMF increased with β (**Fig. 3**). We attribute the poor performance near β=0 to the difficulty of predicting combination effects of drugs that are separated by large distances on the seed network. The adjacency matrix for a regular lattice is banded, with the unknown values comprising a contiguous block. Performance improves for values of β near ½, where the small-world property emerges, and peaks at β=1, the Erdős-Réyni network. Whereas the small-world Watts-Strogatz graph provides a short path between any pair of nodes, the Erdős-Réyni graph contains multiple paths, each carrying evidence for the value of the inferred combination efficacy.

**Figure 3.**
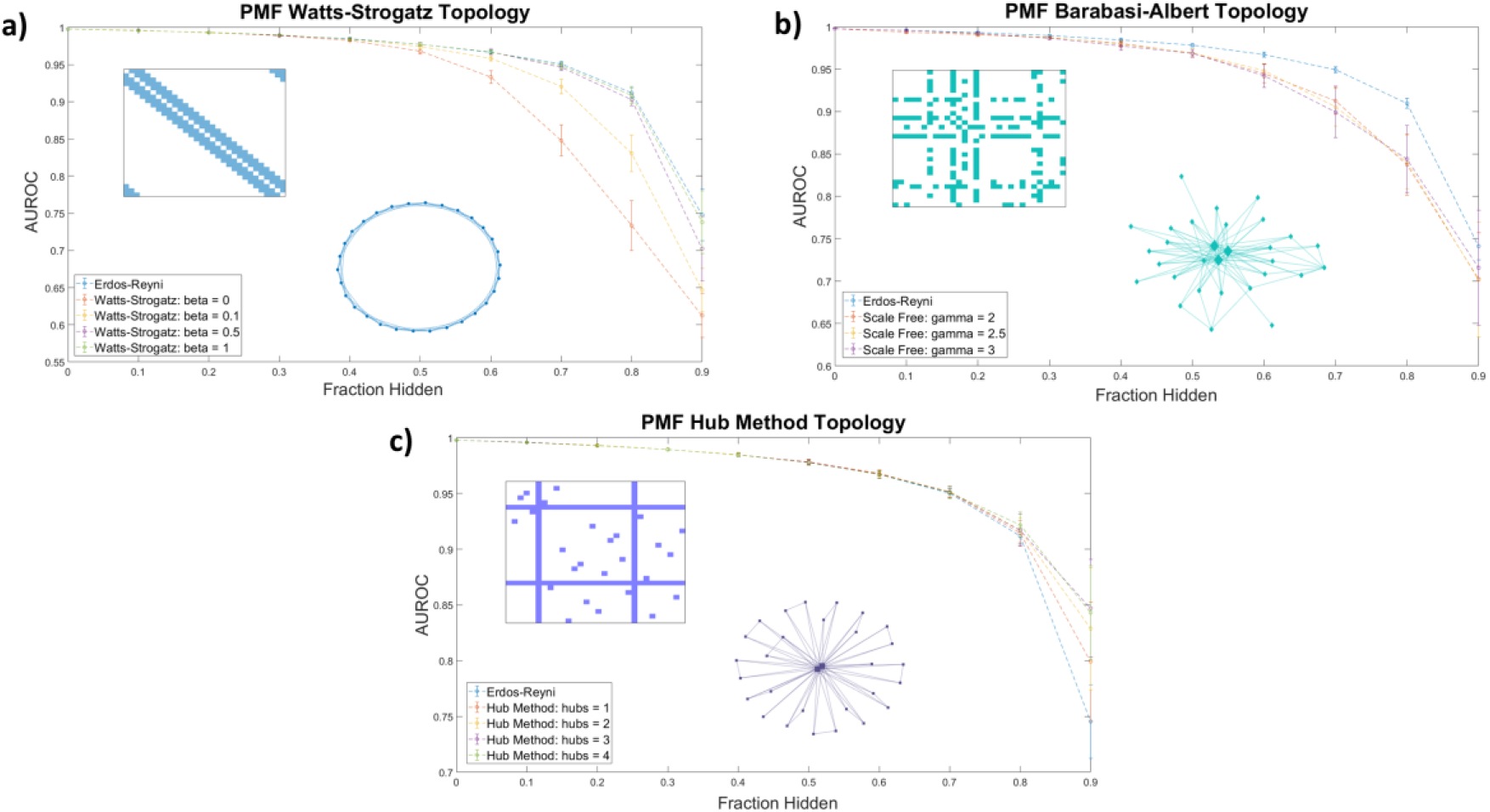
The AUROC of PMF in identifying efficacious combinations as the fraction of the data hidden increases is measured for a) Watts-Strogatz graphs, b) Barabasi-Albert scale-free graphs, and c) graphs generated by the Hub Method. Included in each plot is a sample of the adjacency matrix and topology of each network. Error bars represent standard deviation over 25 repeated trials at the same value of β. **a)** Watts-Strogatz graphs with varying β. When β is near zero, each drug has *k* connections with its nearest neighbors in a lattice structure, and the model performs worse than reproducing from an Erdős-Réyni distribution of equivalent size. As β approaches 1 and the degree distribution of the graph converges to a similar Poisson Distribution of an Erdős-Réyni graph, the accuracy of the predictions begins to approach the level of accuracy seen with purely random topologies. **b)** Scale-free seed networks perform similarly, regardless of scaling exponent. On aggregate, scale-free graphs perform similarly for all values of γ, and slightly underperform compared to Erdős-Réyni topologies. **c)** Graphs generated using the hub method with random hubs produce more accurate predictions than other graph types. When most data are hidden, error and standard deviation of the prediction decrease as the number of hubs increases.

Many real-world networks do not follow a binomial or Poisson degree distribution, and instead follow a power law or scale-free distribution. In a scale-free network, the probability that a node has *k* edges is proportional to *k^-γ^*, where γ is a scaling exponent between 2 and 3. We explored whether a scale-free distribution in the input data influences the accuracy of the prediction. Using the hidden parameter model [19–21], we generated scale-free seed networks for training PMF. The method performed equally well for scale-free distributions for all values of γ, and predicted unknown values with accuracy comparable to the Watts-Strogatz method (**Fig. 3**). Designing a combination screen using the above-described graph topologies may not be experimentally convenient; instead, screeners are more likely to select a few well-known compounds and test them in combination with other compounds in a large library. The adjacency matrix of the seed graph in this approach has several rows/columns in which every value is known, while the large majority of have few or no known values (**Fig. 3c**). The corresponding graph has several fully connected hubs, with the remaining nodes having very few connections.

We explored the accuracy of PMF by using a hub method construction defined as follows: First, we ensured that every node had exactly one connection. Then, we selected nodes at random to be hubs, and ensured that each hub was fully connected. Finally, the remaining edges were randomly assigned in an Erdős-Réyni random fashion. We found the PMF performed stronger on hub method topologies than random Erdős-Réyni topologies when more than 80% of the network was hidden and the graph was sparse. Moreover, when training data was arranged in a hub model topology, the performance of PMF increased as the number of hubs increased (**Fig. 3**).

Thus, we found that the specific seed topology of the training data did not greatly affect the accuracy of the prediction in identifying synergistic drugs if the topology was a random Erdős-Réyni graph or had a binomial degree distribution, such as the Watts-Strogatz for large β. However, PMF did perform worse when edges were evenly distributed following the Watts-Strogatz model for small values of β or when edges were distributed following a scale-free distribution. Moreover, we found that PMF was more accurate under hub topologies mirroring real drug combination assays when more than 80% of the network was hidden, which is exactly the region of interest if we want to test as few combinations as possible.

### PMF as a tool to guide combination screening

In vitro phenotypic-based screens have several benefits for drug discovery, such as not needing to know the molecular target of a disease and being less restricted by hypotheses [22]. However, throughput can be low in such assays, and increasing the number of compounds to be screened causes experimental effort and cost to rise exponentially. PMF may help combat this issue by guiding combination screens through iterative prediction and testing in an active learning scheme.

We simulated PMF being used in an active learning experimental design as follows. First, we created a random Erdős-Réyni graph topology with 10% of the total combinations known. Then, we used PMF to reproduce the entire combination efficacy matrix and identified the top 5% greatest efficacies as predicted by PMF. We then “tested” these identified efficacious combinations by adding the actual values of the efficacies to the list of known combinations, and then repeated the procedure to discover the next 5%, until the entire matrix is recovered. PMF-guided screens identified efficacious combinations much more efficiently than naïve random tests (**Fig. 4**). In our simulated experiment, PMF identified efficacious combinations at three times the rate of random choice and identified as much as 95% of all highly efficacious combinations while only testing 50% of all available combinations. This finding was consistent across cell lines and was not sensitive to the details of the starting point. Our results suggest that screeners may be able to test a small number of relevant combinations of direct interest and obtain the remaining synergistic combinations following a PMF-guided design. Future studies could fruitfully explore this issue further by optimizing PMF-guided screens as well as investigating its accuracy applied in a physical assay experiment.

**Figure 4.**
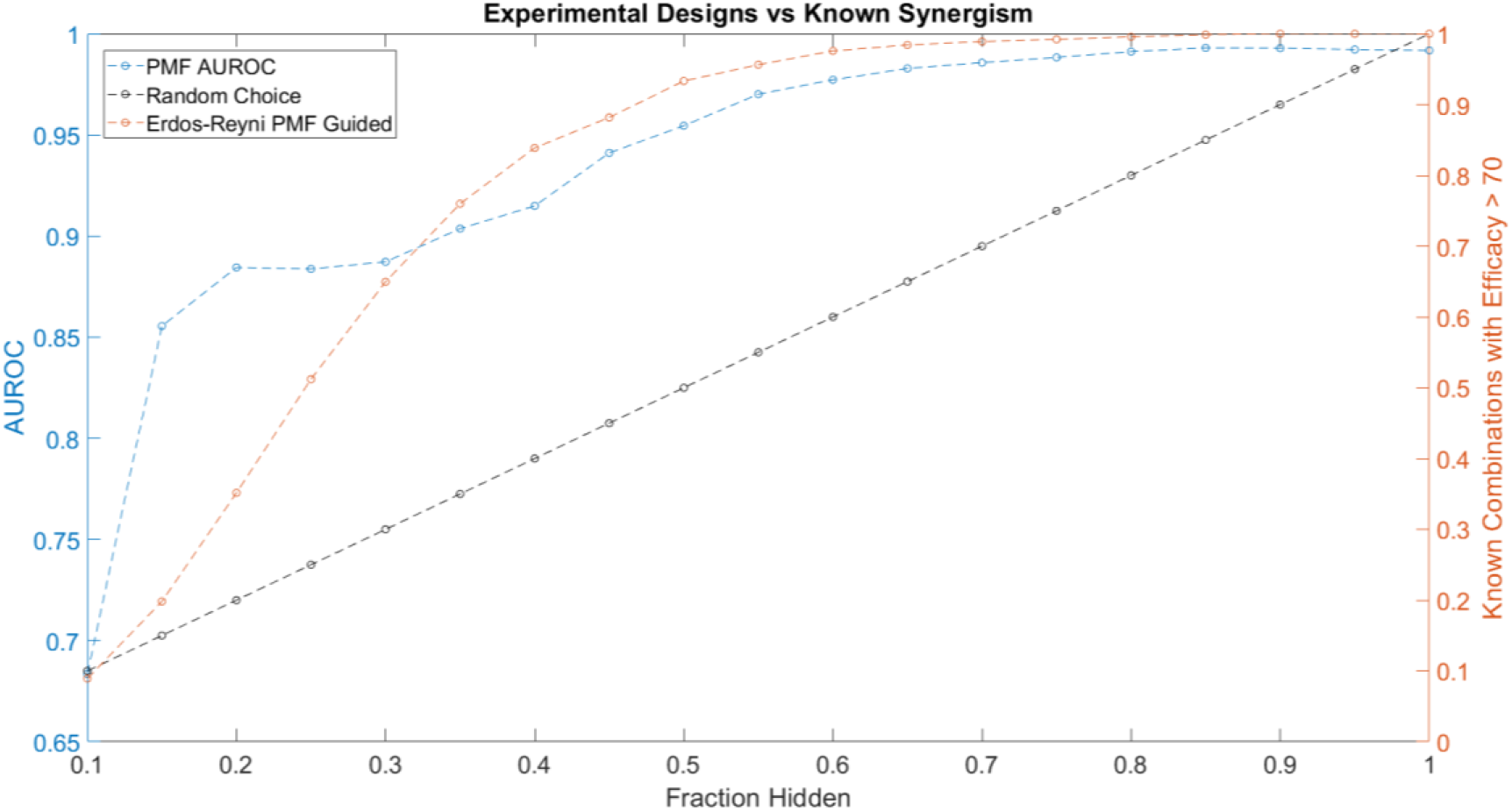
The performance of PMF in a proposed experimental design to predict drug combinations with efficacy greater than 70 is plotted in orange and is compared against random choice plotted in black. Both the AUC of PMF’s predictions in blue as well as the percentage of known efficacious drugs with efficacy greater than 70 are plotted against the known fraction of the drug-drug efficacy matrix. The experiment following random choice takes a random sample of the graph, resulting in a linear relationship between the amount of the drug-synergy matrix known and the amount of known synergistic drugs. As the procedure described above is repeated, PMF identifies more than 95% of the most efficacious drugs while only knowing 50% of the full drug efficacy matrix, much greater than random choice.

Throughout the simulated experiment, we monitored the performance of PMF as measured by AUROC (**Fig. 4**). The dip in AUROC observed around the fourth step of the simulated experiment may be due to bias introduced by the active learning. Efficacious combinations are not uniformly distributed across all drugs, and indeed a small subset of drugs is likely to contribute to many of the efficacious combinations. As the experiment progresses, PMF preferentially selects combinations from an efficacious minority of the nodes, mirroring the construction of a scale-free graph. PMF performs worse on scale-free graphs compared to Erdős-Réyni graphs (**Fig. 3**), causing the accuracy to decrease as nodes are preferentially tested, and then increase as these nodes are saturated and the rest of the matrix is tested. Future studies might investigate ways to counteract this drop in error by using a more complex method than simply testing the top 5% most efficacious combinations as predicted by PMF.

### PMF predicts efficacy, but not synergy

The desired output of most phenotypic combination screens is an efficacious and non-toxic combination; however, *de novo* development of combination therapeutics will benefit from identifying synergistic drug combinations, whether or not they are efficacious. For example, two drugs that individually have no efficacy may have a moderate effect in combination. Although such a combination may not be clinically useful, it carries structure and pathway information that may serve as the starting point for rational development of combination therapeutics. PMF was able to recover missing values much less accurately when predicting synergy rather than efficacy. Just as with efficacies (**Fig. 1**), PMF recovered training data to arbitrary precision (**Fig. 5a**), but it did not recover test data well, unless it had a sufficiently large training set (i.e., small fraction of data hidden) (**Fig. 5b-c**). While the accuracy of PMF on predicting synergy was much weaker than PMF predicting efficacy, we still found that the model is robust and performed well in cases where 50-70% of the total matrix was known.

**Figure 5.**
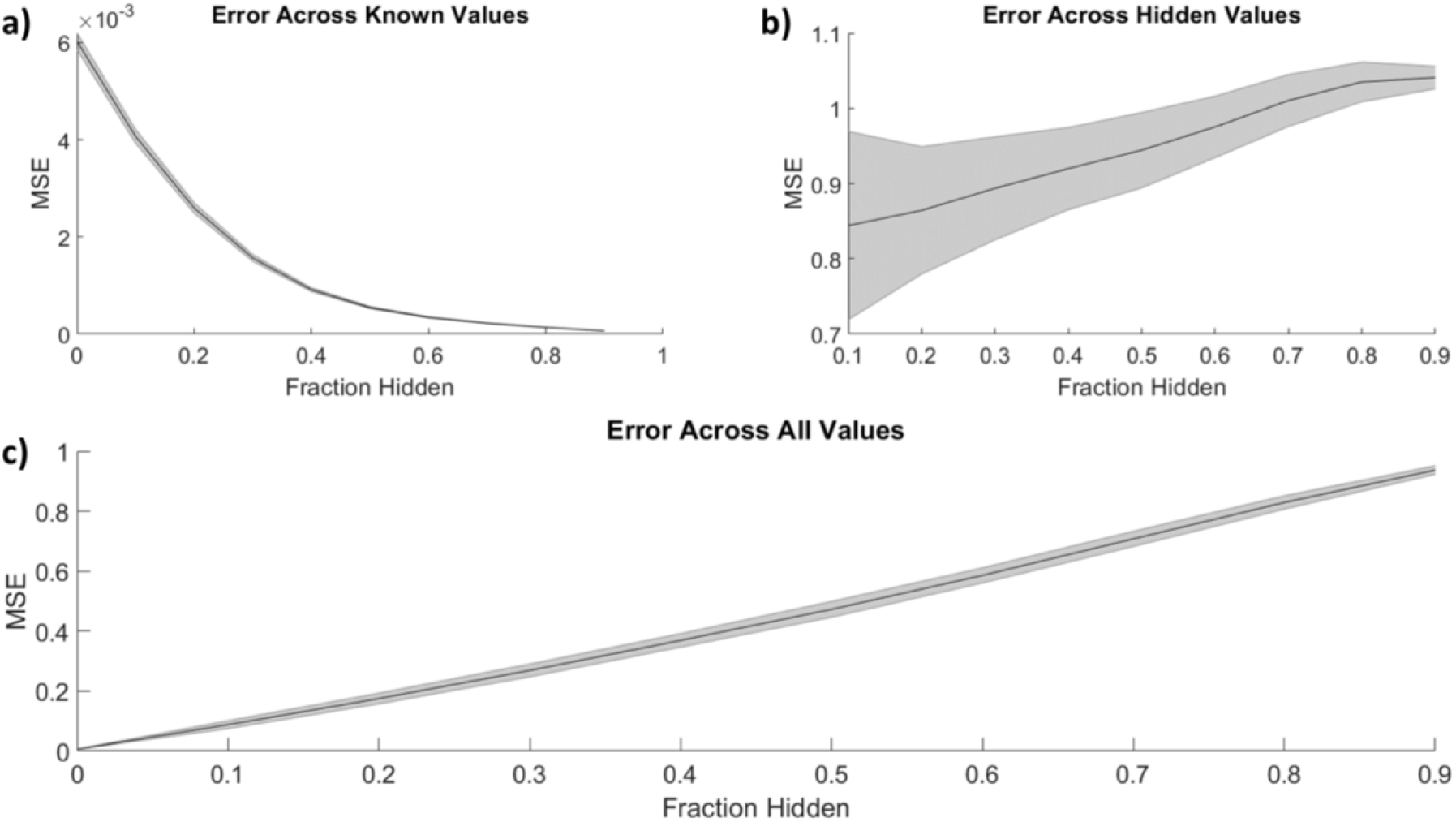
Similar to **Fig. 1**, the mean-squared error of PMF in recovering combination synergy of a) known, b) hidden, and c) all elements is plotted against the fraction of hidden data. In all panels, the shaded area represents the standard deviation of the mean-squared error over 25 trials across all cell lines. Once again, PMF recovers all known data to arbitrary precision. PMF performs with much less accuracy when predicting ComboScores rather than efficacies (**Fig. 1**). Error is much greater and more uncertain overall in hidden indices and thus across all indices.

This stark decrease in accuracy and predictive power may result from the lopsided definition of synergy. The ComboScore of each drug combination represents the difference between the observed effect of the combination and the expected effect assuming each drug acts independently. Because the upper bound on efficacy is the same for individual drugs and combinations, a combination of highly efficacious drugs cannot have a high ComboScore, even if it has optimal efficacy. Similarly, combinations with identical efficacies may have different ComboScores, depending on the efficacies of the individual drugs used in the combinations. Thus, a low ComboScore reveals nothing about the efficacy of the combination, but a high ComboScore indicates the combination’s component drugs individually have low efficacy (**Suppl. Fig. 2**). As ComboScores are calculated from individual and combination efficacies, one can still use PMF to predict combination efficacies, and use these to calculate ComboScores.

## Conclusion

We have shown that PMF can accurately impute missing values into the drug combination efficacy matrix for a screen, and that the performance of PMF does not depend on the efficacies of the drugs being tested. We further showed that PMF performs best when the input drug combination network has an Erdős-Réyni topology. Finally, we used simulated experiments to demonstrate that alternating PMF inference with experiments can efficiently identify the most efficacious two-drug combinations in a phenotypic screen.

There have been many other attempts at predicting the effects of drug combinations, and those that perform best include additional data, such as chemical structures, target profiles, or OMICS data[23–25]. Our method is simpler by comparison, but it provides a baseline of performance against which more complicated prediction methods may be assessed. Indeed, not relying on additional information endows our method with flexibility: Instead of predicting the effects of combinations of drugs, it can be used to predict the effects of combinations of combinations, and we have no reason to believe that it will perform worse on unannotated compounds. On the contrary, our method may contribute to identifying mechanisms of action for novel compounds. The very ability of PMF to predict efficacies of combinations points to hidden mechanistic similarities within the set of compounds. By interpreting the PMF in terms of underlying biochemistry, we may gain insight into the nature of disease.

## Acknowledgements

This work was supported by the National Institutes of Health through grants UL1TR001857 and R56AG059612.

