## Supplementary material for "Predicting the effects of drug combinations using probabilistic matrix factorization": Suppl. Fig.

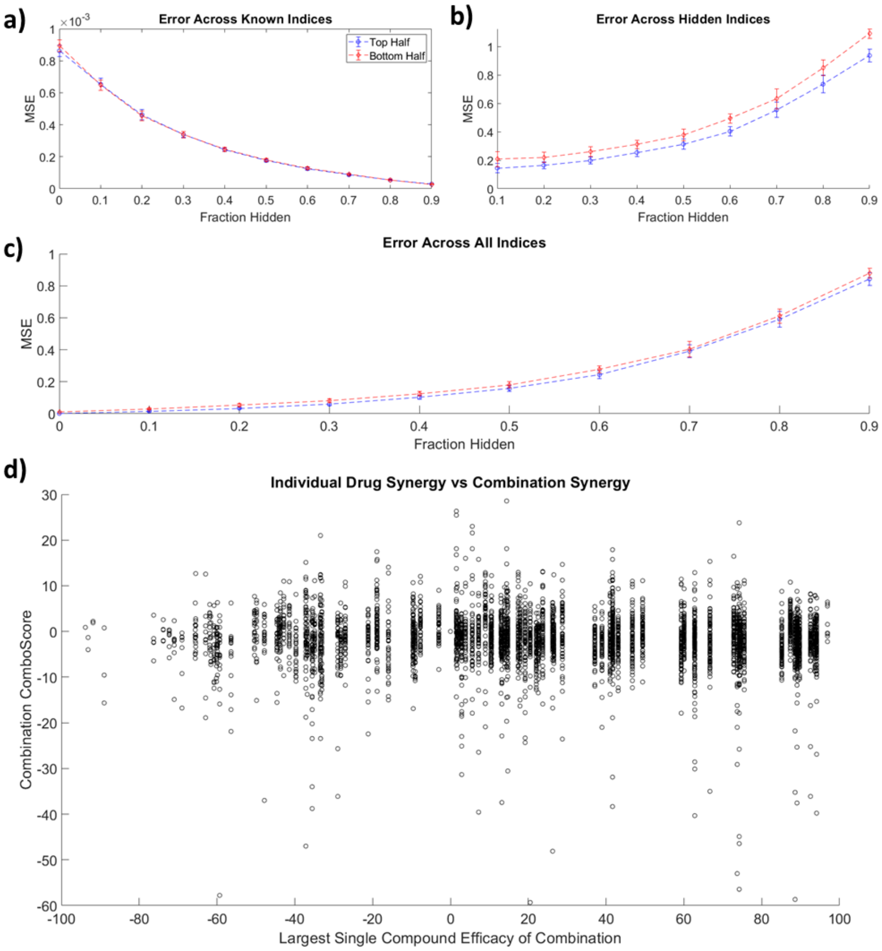


**Supplementary Figure 1.** Similarly to Figure 1, the mean-squared error of PMF in recovering values of a) known, b) hidden, and c) all elements on the 786-0 cell line, when training data is biased towards only individually efficacious or inefficacious drugs is plotted against the fraction of hidden data. The error across known indices remains small and identical regardless of how the training data is biased. PMF is slightly less accurate at predicting unknown data when it is biased towards weakly efficacious drugs than highly efficacious drugs. d) Combination efficacies are plotted against the efficacy of the individual drugs, showing that individually efficacious compounds may not lead to efficacious combinations.

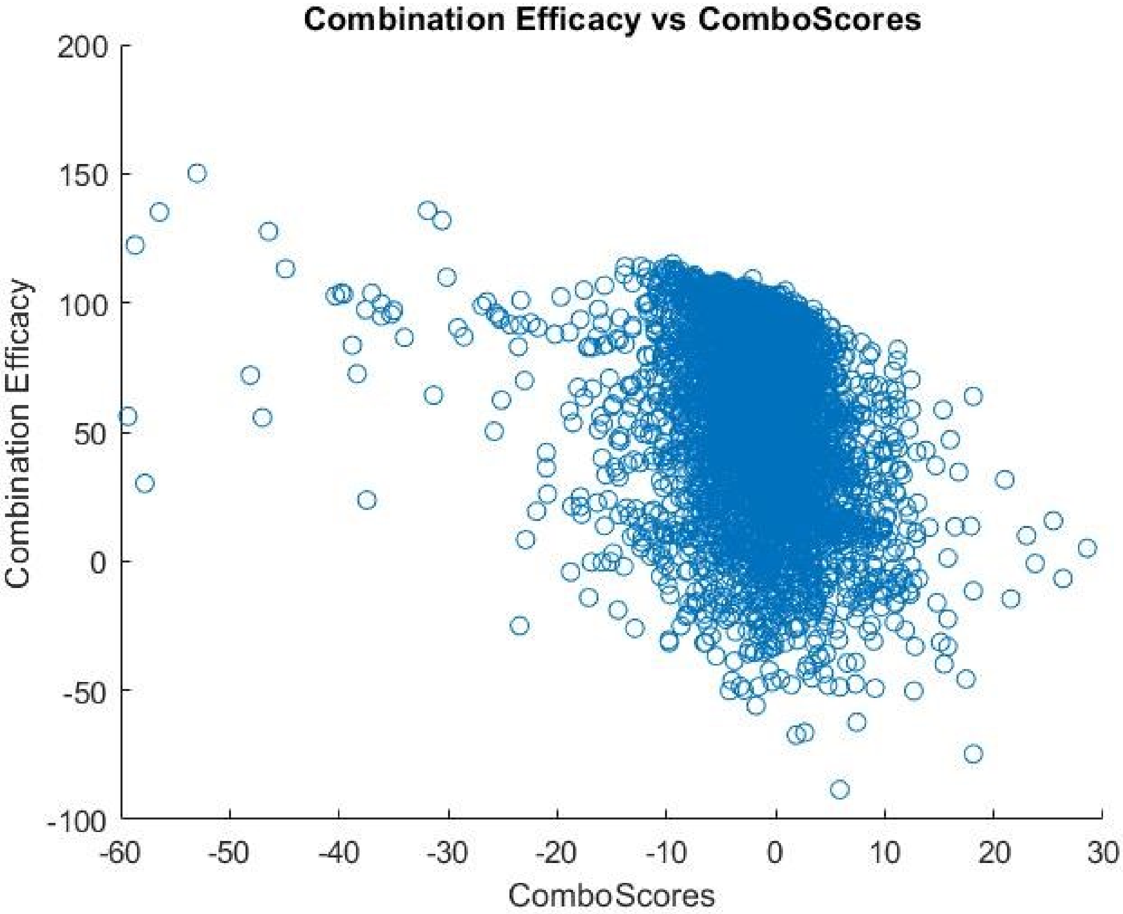


**Supplementary Figure 2.** The ComboScore of each drug Combination is plotted against the efficacy of the combination. Clearly, there is no strong correlation between efficacy and synergy, making synergy a much more difficult and abstracted concept to measure.
